# rMSIcleanup: An open-source tool for matrix-related peak annotation in mass spectrometry imaging and its application to silver-assisted laser desorption/ionization

**DOI:** 10.1101/2019.12.20.884957

**Authors:** Gerard Baquer, LLuc Sementé, María García-Altares, Young Jin Lee, Pierre Chaurand, Xavier Correig, Pere Ràfols

## Abstract

Mass spectrometry imaging (MSI) has become a mature, widespread analytical technique to perform non-targeted spatial metabolomics. However, the compounds used to promote desorption and ionization of the analyte during acquisition cause spectral interferences in the low mass range that hinder downstream data processing in metabolomics applications. Thus, it is advisable to annotate and remove matrix-related peaks to reduce the number of redundant and non-biologically-relevant variables in the dataset. We have developed rMSIcleanup, an open-source R package to annotate and remove matrix-related signals based on its chemical formula and the spatial distribution of its ions. To validate the annotation method, rMSIcleanup was challenged with several images acquired using silver-assisted laser desorption ionization MSI (AgLDI MSI). The algorithm was able to correctly classify *m/z* signals related to silver clusters. Visual exploration of the data using Principal Component Analysis (PCA) demonstrated that annotation and removal of matrix-related signals improved spectral data post-processing. The results highlight the need for including matrix-related peak annotation tools such as rMSIcleanup in MSI workflows.

**Resources availability:** The R package presented in this publication is freely available under the terms of the GNU General Public License v3.0 at https://github.com/gbaquer/rMSIcleanup. The datasets used in the experiments can be accessed upon request to the corresponding author.

## 1. Introduction

Mass spectrometry imaging (MSI) is a label-free technology that allows to obtain molecular and spatial information from intact tissue sections [1]. MSI has been gradually adopted for spatial-resolved metabolomics and it has been regarded as a potential tool for understanding the mechanisms underlying complex diseases such as cancer or diabetes [2]. However, the conventional organic matrices used in Matrix-Assisted Laser Desorption Ionization (MALDI) produce spectral signals that interfere in the low *m/z* range. This is an issue particularly in metabolomics which analyses low molecular weight compounds, so mass spectrometers are set to acquire within the *m/z* range where MALDI matrices exhibit most MS signals. This seriously hampers downstream metabolomics data processing [3,4], as the matrix introduces noise, redundant variables, and variables with no biological meaning into the complex MSI datasets.

Several alternatives to the common organic matrices have been proposed to deal with exogenous contamination caused by matrix ion signals. Nanomaterials or metal layer deposition methods, for instance, dramatically reduce the number of signals related to the LDI promoting material in the low *m/z* range. Some examples are graphene oxide, silicon or metals such as gold, platinum or silver [5–8]. Nevertheless, even when these alternatives are used and the number of peaks related to the LDI promoting material is reduced, there is still a need to annotate them in order to reduce spectral complexity and distinguish exogenous from endogenous compounds, especially in untargeted applications.

To tackle the issue of annotating MS signals related to the LDI-promoting material several software-based solutions have been proposed. A simple approach consists of acquiring a reference area outside the sample during the MSI experiment. Under the assumption that only matrix-related peaks will be recorded, the peaks found in the outside area are then subtracted from the tissue spectrum. Given its simplicity, some variation of this procedure has been adopted by many researchers in their workflows. Expanding on this idea, Fonville et al. [9] presented a method that relies on the hypothesis that matrix-related peaks will correlate positively to a set of reference peaks outside the tissue region while endogenous peaks will correlate negatively. However, this approach has three main limitations. Firstly, due to ion suppression [10] and the formation of matrix adducts with endogenous compounds, the matrix-related peaks outside and inside the tissue region might differ. Additionally, endogenous molecules that are delocalized during the matrix application process can be misclassified as matrix-related. Finally, the method cannot distinguish a given matrix-related MS peak from an isobaric or overlapping endogenous MS peak. Thus, simplified approaches to annotate matrix-related signals are not suitable for untargeted applications such as spatial metabolomics. Recent work by Ovchinnikova et al. [11] takes a more comprehensive approach in defining three automated algorithms for off-sample ion classification. Their methods have proved to perform well when trained and validated against a “gold standard set” of ion images manually annotated by experts. However, their focus is not specifically on matrix-related peaks, but on the annotation of signals that exhibit a spatial distribution with high concentrations outside of the tissue region. For this reason, these methods focus on classifying each ion image separately as “on-sample” or “off-sample” and do not exploit relevant information such as the identity of the ion, adduct type, matrix type, etc. Additionally, since they are based in machine and deep learning methods they inherently suffer from the black box problem given that annotation results cannot be traced back and easily justified.

To solve these limitations we propose a new algorithm that relies not only on the ion images but also on the chemical information of the LDI promoting material used. The algorithm also incorporates an overlapping peak detection feature to prevent misclassification of overlapped or isobaric ions. The presented algorithm is implemented in an open-source R package freely available to facilitate its use. Additionally, the package generates a visual report to transparently justify each annotation.

In order to validate and optimize the proposed method, we opted for a well-understood LDI promoting material such as silver. The use of silver nanolayers for MSI (AgLDI MSI) has been steadily growing in recent years [6,12–17]. The characteristic isotopic pattern of silver (^107^*Ag* and ^109^*Ag*, 51.84% and 48.16% abundance, respectively), as well as its well-known ionization and adduct formation allow to define a list of possible and not-possible silver-related peaks of a typical AgLDI MSI experiment. This set of possible and not-possible peaks is used as a validation list to assess the performance of the classification algorithm. A total of 14 MSI datasets acquired with an Ag-sputtered nanolayer from three different laboratories, were used for validation.

## 2. Materials & Methods

Table 1 summarizes the main processing parameters for each of the 14 datasets used in this study. Datasets 1-10 were acquired in our lab and the materials, sample preparation and MSI acquisition parameters are described here. In order to overcome lab-specific bias in our study, four additional datasets were provided by collaborating laboratories. For further details about the materials, sample preparation and MSI acquisition of these datasets, refer to the original publications of Dataset 11 [18], Dataset 12 [14] and Datasets 13 and 14 [6].

**Table 1.** List of the 14 AgLDI MSI datasets used for validation. Sample type, sample preparation and LDI-MSI acquisition parameters. Datasets from 1-10 were acquired in-house. Datasets 11-14 were provided by external laboratories.

| No. | Species | Tissue type | Ag deposition system and estimated layer thickness | Lateral Res. | m/z range | Mass spectrometer | Acq. Mode | Ref. |
| --- | --- | --- | --- | --- | --- | --- | --- | --- |
| 1 | Mouse | Pancreas | ATC Orion 8-HV Sputtering system, 5 nm | 30 $\mu m$ | 70-1200 | Bruker ultrafleXtreme™ MALDI-TOF/TOF | Positive / Profile | - |
| 2 | Mouse | Pancreas | ATC Orion 8-HV Sputtering system, 5 nm | 30 $\mu m$ | 70-1200 | Bruker ultrafleXtreme™ MALDI-TOF/TOF | Positive / Profile | - |
| 3 | Mouse | Kidney | ATC Orion 8-HV Sputtering system, 5 nm | 100 $\mu m$ | 70-1200 | Bruker ultrafleXtreme™ MALDI-TOF/TOF | Positive / Profile | - |
| 4 | Mouse | Brain | ATC Orion 8-HV Sputtering system, 5 nm | 80 $\mu m$ | 70-1200 | Bruker ultrafleXtreme™ MALDI-TOF/TOF | Positive / Profile | - |
| 5 | Mouse | Brain | ATC Orion 8-HV Sputtering system, 5 nm | 80 $\mu m$ | 70-1200 | Bruker ultrafleXtreme™ MALDI-TOF/TOF | Positive / Profile | - |
| 6 | Mouse | Brain | ATC Orion 8-HV Sputtering system, 5 nm | 80 $\mu m$ | 70-1200 | Bruker ultrafleXtreme™ MALDI-TOF/TOF | Positive / Profile | - |
| 7 | Mouse | Brain | ATC Orion 8-HV Sputtering system, 5 nm | 80 $\mu m$ | 80-1000 | Bruker ultrafleXtreme™ MALDI-TOF/TOF | Positive / Profile | - |
| 8 | Mouse | Brain | ATC Orion 8-HV Sputtering system, 5 nm | 80 $\mu m$ | 80-1000 | Bruker ultrafleXtreme™ MALDI-TOF/TOF | Positive / Profile | - |
| 9 | Mouse | Brain | ATC Orion 8-HV Sputtering system, 5 nm | 80 $\mu m$ | 80-1000 | Bruker ultrafleXtreme™ MALDI-TOF/TOF | Positive / Profile | - |
| 10 | Mouse | Brain | ATC Orion 8-HV Sputtering system, 5 nm | 80 $\mu m$ | 80-1000 | Bruker ultrafleXtreme™ MALDI-TOF/TOF | Positive / Profile | - |
| 11 | Mouse | Brain | Cressington Sputter Coater, 23 ± 2 nm | 75 $\mu m$ | 100-1100 | Bruker ultrafleXtreme™ MALDI-TOF/TOF | Positive / Profile | [18] |
| 12 | *Homo sapiens sapiens* | Fingermark | Cressington Sputter Coater, 14 ± 2 nm | 75 $\mu m$ | 100-1100 | Bruker ultrafleXtreme™ MALDI-TOF/TOF | Positive / Profile | [14] |
| 13 | B73 inbred corn | Root | Cressington 108Auto, 5s | 10 $\mu m$ | 50-970 | Thermo Finnigan™ MALDI-LTQ-Orbitrap Discovery | Positive / Centroid | [6] |
| 14 | B73 inbred corn | Root | Cressington 108Auto, 5s | 10 $\mu m$ | 50-900 | Thermo Finnigan™ MALDI-LTQ-Orbitrap Discovery | Negative / Centroid | [6] |

### 2.1. Materials

For the samples acquired by our group, indium tin oxide (ITO)-coated glass slides were obtained from Bruker Daltonics (Bremen, Germany). The silver-target (purity grade > 99.99%) used for sputtering was acquired from Kurt J. Lesker Company (Hastings, England).

### 2.2. Sample preparation

All the samples acquired by our group were obtained from mice and provided by the animal facility at the Faculty of Medicine and Health Sciences of the University Rovira i Virgili. All tissues were snap-frozen at −80°C after collection and kept at this temperature during shipping and storing until MSI acquisition.

The tissues were sectioned with a Leica CM-1950 cryostat (Leica Biosystems Nussloch GmbH) located at the Centre for Omics Sciences (COS) of the University Rovira i Virgili into 10 *μm* sections. Tissue sections were mounted on ITO coated slides by directly placing the glass slide at ambient temperature onto the section.

The sputtering system ATC Orion 8-HV (AJA International, N. Scituate, MA, USA) was used to deposit a silver nanolayer onto each tissue section. An argon atmosphere with a pressure of 30 mTorr was used to create the plasma in the gun. The working distance of the plate was set to 35 mm. The sputtering conditions were ambient temperature using DC mode at 100W for 10s. With these parameters, an Ag layer thickness of roughly 5nm was obtained. The deposition times were short to prevent the substrate temperature from increasing excessively and, consequently, degrading metabolites.

### 2.3. LDI-MS acquisition

A MALDI TOF/TOF ultrafleXtreme instrument with SmartBeam II Nd:YAG/355 nm laser from Bruker Daltonics available at COS was used for MSI acquisition. Acquisitions were carried out by operating the laser at 2 kHz and collecting a total of 500 shots per pixel.

The TOF spectrometer was operated in positive ion, reflectron mode, in *m/z* ranges according to Table 1. The spectrometer was calibrated prior to MSI data acquisition using 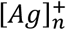 cluster peaks as internal reference masses.

### 2.4. MSI data processing

The raw spectral data of each MSI dataset was exported to the imZML data format [19] in profile mode. The software rMSIproc [20] was used to process the data and generate a peak matrix in centroid mode. The default processing parameters were used. The Signal-to-Noise Ratio (SNR) threshold was set to 5 and the Savitzky-Golay smoothing had a kernel size of 7. Peaks appearing in less than 5% of the pixels were filtered out. Peaks within a window of 6 data-points or scans were binned together as the same mass peak. Mass spectra were re-calibrated using the Ag reference peaks as reference masses [21].

Datasets 13 and 14 were acquired in centroid mode with an Orbitrap mass spectrometer. These datasets were directly submitted to the binning process of rMSIproc [21] to conform to the peak matrix format.

No data normalization was performed. Data were visualized and explored using rMSI [22].

## 3. Algorithm description

### 3.1. Input and output format

The matrix-related annotation algorithm takes the peak matrix in centroid mode and the processed spectral data in profile mode as input. The user must also provide the chemical formulae of the matrix applied and a list of possible adducts and neutral losses to consider.

The algorithm produces a vector containing the similarity scores that indicate the likelihood of each mass in the input image being a matrix-related ion. The package also provides an informative visual report for the user to understand the justification behind the classification. Supplementary Figures S1-S4 show examples of the visual report.

### 3.2. In-silico cluster & adduct calculation

The theoretical mass and relative isotopic pattern intensities of all possible matrix-related clusters are calculated using the open-source package enviPat [23], a fast and memory-efficient algorithm to compute theoretical isotope patterns.

For each theoretical cluster *t*_*i*_ its experimental counterpart *e*_*i*_ is obtained from the mean spectra of the dataset. The experimental masses closest to the theoretical ones within a given tolerance specified by the user are used. The theoretical clusters will then be matched against their experimental counterparts and their presence in the experimental dataset assessed using two similarity metrics.

### 3.3. Similarity metrics

The similarity between each theoretical matrix-related cluster and experimental clusters is assessed using two similarity scores according to equation 1.

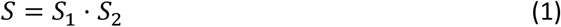

where *S* is the total similarity score, *S*_1_ is the cluster spectral similarity and *S*_2_ is the intra-cluster morphological similarity. Both similarity scores range from 0 to 1.

The cluster spectral similarity score *S*_1,*i*_ for theoretical cluster *t*_*i*_ determines the degree of similarity between the scaled intensity vectors of intensities 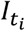 and 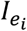 and it is computed according to equation 2.

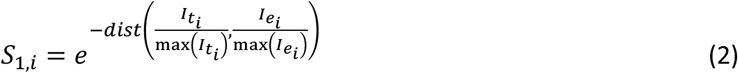

where *dist*(*a, b*) is the distance function chosen by the user (Euclidean distance by default), 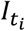 is the vector of intensities of the theoretical cluster *t*_*i*_ and 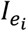 is the vector of intensities of experimental cluster *e*_*i*_. Experimental cluster *e*_*i*_ is determined by accessing the element in the peak matrix with the mass closest to that corresponding to *t*_*i*_ within a given tolerance. In plain terms, *S*_1_ is a decaying exponential function of the distance between the intensity scaled intensity vectors 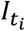 and 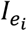.

The intra-cluster morphological similarity *S*_2,*i*_ returns the degree of similarity between the spatial distributions of the ions conforming the experimental cluster *e*_*i*_. Ions with a high spatial correlation are more likely to belong to the same cluster. This metric is computed using equation 3.

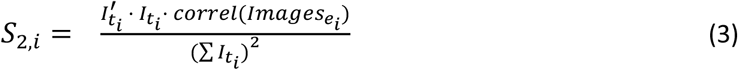

where 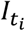 is the intensity vector of the theoretical cluster *t*_*i*_, *correl*(*A*) is the correlation function specified by the user (Pearson correlation by default) and 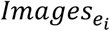 is the set of images corresponding to each ion in the experimental cluster *e*_*i*_. In plain terms, *S*_2_ is the weighted mean across both directions of the correlation matrix between each ion image in *e*_*i*_.

### 3.4. Overlapping peak detection

Insufficient resolving power leads to overlapped MS signals, which can be a severe problem in matrix-related peak annotation as they can lead to a greater number of misclassified peaks. This is a particularly limiting issue in lower resolution spectrometers such as some TOFs in contrast to higher resolution analysers such as Orbitrap or FTICR [24]. An additional problem with the same effect is the intrinsic inability of mass spectrometry to distinguish between isobaric species. In order to cope with these issues, we propose an overlapping detection algorithm capable of determining if a given MS signal corresponds to more than one overlapped ion peaks.

The overlapping detection algorithm is only executed in those clusters that report S1 and S2 scores under a threshold specified by the user. Before concluding that the cluster is not present, the algorithm determines whether the low similarity metrics could be attributed to the presence of overlapped signals.

The algorithm is based on the operating principle of bisecting k-means [25]. All the ions in an experimental cluster *e*_*i*_ are split into two subgroups (*e*_*i*:1_ and *e*_*i*:2_) based on the correlation of their spatial distributions using k-means. For each subgroup of ions the similarity metrics S1 and S2 are recomputed. If the S1 and S2 scores of a given subgroup surpass the specified threshold, all ions in the subgroup are tagged as matrix-related. The remaining ions in *e*_*i*_ are tagged as matrix-related but suffering from overlapping, and the overlapping detection algorithm terminates. If instead, none of the subgroups obtains an S1 and S2 above the threshold, the process of splitting into two subgroups by k-means and recomputing the similarity scores is repeated for both *e*_*i*:1_ and *e*_*i*:2_. This bisection of the ions in *e*_*i*_ is repeated iteratively until a subgroup obtains S1 and S2 scores above the threshold. To prevent overfitting, the iterative process will also stop when the number of peaks contained by the biggest subgroup becomes smaller than half the amount of peaks in *e*_*i*_. In such event, it is concluded that there are no overlapped peaks and all ions in the experimental cluster *e*_*i*_ are tagged as not-matrix-related. To sum, overlapped MS signals will be detected and distinguished from the rest of the ions in the cluster based on the dissimilarity of their spatial distributions.

## 4. Results

### 4.1. Algorithm validation with AgLDI MSI

In order to validate and optimize the algorithm, we opted to use sample tissues covered by silver nanoparticles, a well-defined and understood LDI promoting material. A total of 14 datasets, from 3 different laboratories, were used. The datasets included several animal tissues, plant tissues and human fingermarks.

The algorithm was challenged with the task of classifying a list of silver-containing compounds and adducts for each dataset. The list includes a “positive class” formed by clusters that should be present in all samples used in this study and a “negative class” containing clusters that should not be present in any of them. This list is referred to as “validation list” and allowed us to assess the performance of the algorithm. An algorithm with a perfect performance should classify all clusters in the “positive class” as matrix-related signals and all clusters in the negative class as not present and thus not-matrix related. This is a common approach in bioinformatics for validating and assessing the performance of a classifier algorithm [26]. Table 2 shows the complete validation list.

**Table 2.**
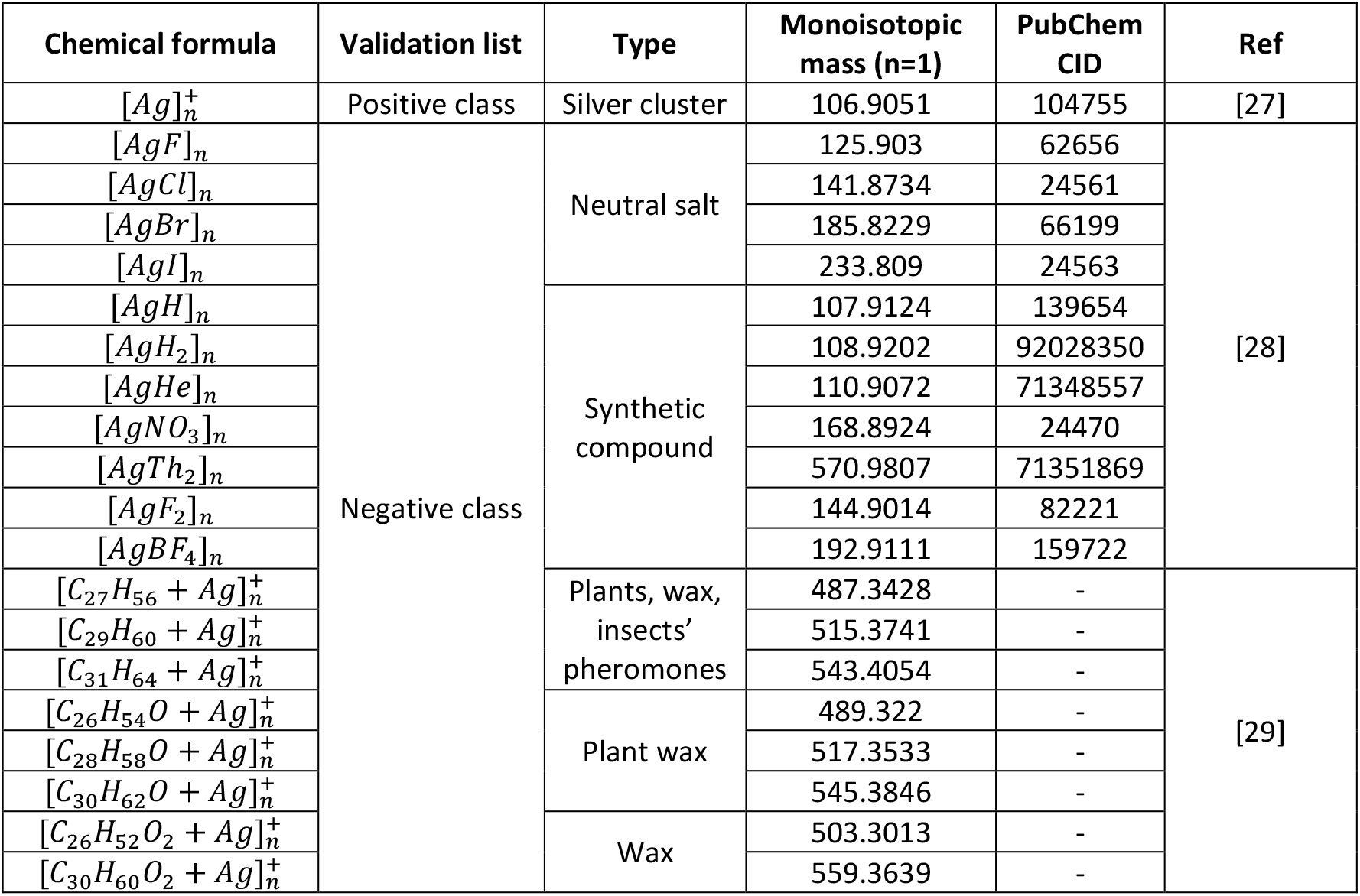
“Validation list” used for validation. The “positive class” consists of silver clusters. The “negative class” consists of neutral silver salts, synthetic silver compounds and silver adducts that are not expected to be found in animal samples. The index n denotes the number of atoms or molecules inside the cluster. The minimum and maximum value of n depend on the monoisotopic mass of the atom or molecule and the mass range of the dataset.

Silver clusters containing up to 60 atoms have been reported to form during silver sputtering [27]. The “positive class” expected to be found in all datasets is therefore formed by all silver clusters within the acquired mass range. For most of the datasets, this includes clusters from 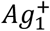 to 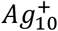.

The “negative class” consists of silver compounds or adducts that should not be present in any of the samples used in this study. Firstly, this list includes various silver neutral salts which cannot be measured using LDI MSI, and some synthetic compounds that are not expected to be present in animal or plant samples [28]. It also includes compounds found in aerial parts of plants, wax and insects (not found in mammal tissues nor in corn root) that have been reported to form adducts with silver in AgLDI MSI applications [29]. For each of these molecules, we also included all clusters within the acquired mass range. These particular molecules and their clusters were selected in an attempt to have a “negative class” covering the full mass range.

### 4.2. Performance of similarity scores

Using the validation list described in section 4.1, we assessed the performance of the similarity scores as a classifier to annotate 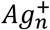-related peaks in AgLDI MSI datasets.

Figure 1 shows the similarity scores obtained for each cluster in Table 2 when searched in all 14 datasets from Table 1. The blue points represent the “positive class” (clusters that should be present) while the red points represent the negative class (clusters that should not be present).

**Fig. 1.**
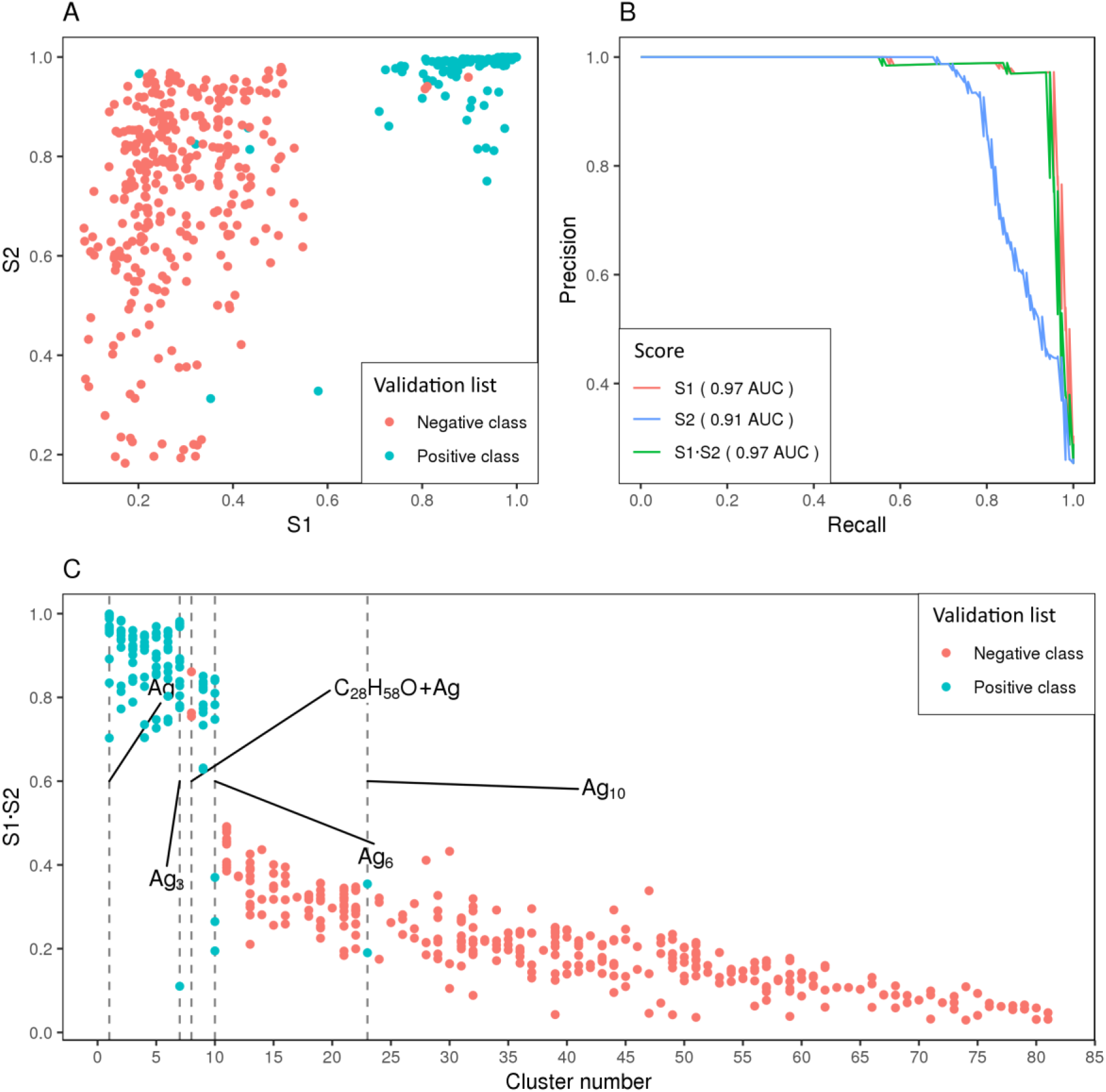
Similarity scores performance (A) Spectral similarity S1 vs. Intra-cluster morphological similarity S2 scatter plot. Each point represents a potential cluster classified by the algorithm. All clusters shown in Table 2 are evaluated for all 14 datasets presented in Table 1. Blue points represent the “positive class” (should be present in the sample) while the red points correspond to the negative class (should not be present in the sample). Most “positive class” points are located in the top right corner well separated from the negative class points. This indicates proper classification power. (B) Precision and recall (PR) curve computed according to Davis et al. 2006 [40]. (C) Similarity score S1·S2 vs. Cluster number. Clusters are arranged in decreasing order of mean similarity score. A clear gap between an S of 0.5 and 0.7 separates the “positive class” from the negative class. Refer to Supplementary Table S1 for a mapping of cluster numbers to cluster chemical formula.

Figure 1A represents the spectral similarity score (*S*1) against the intra-cluster similarity score (S2) of each of these clusters. The “positive class” is clearly separated on the top right corner (high *S*1 and high *S*2).

To evaluate the classifying performance of the two similarity metrics we use the Precision vs. Recall (PR) curve [26]. The precision is defined as the ratio between the number of clusters in the “positive class” classified as matrix-related (i.e. true positives) and the total number of clusters classified as matrix-related (i.e. true positives + false positives). The recall, on the other hand, is the ratio between the number of clusters in the “positive class” classified as matrix-related (i.e. true positives) and the total number of clusters in the “positive class” (i.e. true positives + false negatives). Figure 1B shows the PR curves for each of the similarity metrics proposed. The areas under the curve (AUC) of 0.97 and 0.91, respectively, show that the spectral similarity score S1 is the best classifier followed by the intra-cluster morphology similarity score *S*2. The product of *S*1 · *S*2 had the same classifying skill as *S*1 with an AUC of 0.97. These results prove that 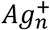-related peaks can be well classified by these two metrics.

*S*1 performs much better than *S*2 as a classifier, and the product of *S*1 · *S*2 matches but does not improve the performance of *S*1 alone. Nevertheless, we still decided to use the product of *S*1 · *S*2 as a classifier in rMSIcleanup instead of using *S*1 alone due to three main reasons. Firstly, the overlapping detection algorithm strongly relies on the morphological similarity of ions and thus depends on *S*2. Moreover, even though we did not find a single instance of a cluster with a high *S*1 score and a low *S*2 score (matching isotopic patterns but unmatching spatial distributions) in any of the samples, we still consider that *S*2 should be present to allow for correct classification should this occur. Finally, *S*2 can be a strong asset in applications other than AgLDI MSI where, due to less distinctive isotopic ratios, the performance of *S*1 as a classifier is diminished.

Figure 1C shows the similarity score S1·S2 obtained by each cluster in all datasets. Clusters are arranged in decreasing order of mean similarity score. Supplementary Table S1 maps the cluster numbers to cluster chemical formula. A clear gap between an *S* of 0.5 and 0.7 separates the “positive class” from the negative one.

Only three false positives (i.e. clusters that should not be present but have a high *S* value) were reported for adduct [C_28_H_58_O + Ag]^+^. An example is shown for Dataset 4 in Supplementary Figure S5. Identification by MS/MS is required to assess if the compound is indeed present in the sample. Nevertheless, the mass error between experimental and theoretical isotopic patterns for this compound was 154 ppm, an error much higher than the expected for this dataset (acquired with a TOF MS analyzer). Therefore, we inferred that the experimental pattern detected is not related to adduct [C_28_H_58_O + Ag]^+^ and this is, in fact, a false positive. In order to reduce the number of false positives, the mass tolerance of the algorithm can be decreased, however, a too strict mass tolerance increases the number of false negatives.

A total of six false negatives (i.e. clusters that should be present but have a low *S* value) were reported for some datasets for clusters *Ag*_3_, *Ag*_6_ and *Ag*_10_. False negatives correspond to clusters for which the majority of peaks in their isotopic pattern were under the SNR threshold, and thus were excluded during pre-processing. In these cases, the few included peaks were not sufficient to reliably annotate the cluster. Supplementary Figure S6 shows the only exception, the *Ag*_6_ cluster in Dataset 12, whose misclassification is not due to intensity problems. In this case, the fingerprint analysed showed highly homogeneous ion images, which impedes the proper operation of the overlapping algorithm and leads to misclassification. Representative examples of correct annotations are shown in Supplementary Figures S7-S8.

As an additional validation, the results were matched against the published annotations of the datasets provided by external laboratories. Dataset 12 contains 60 identifications by MS/MS [14]. Dataset 13 contains 4 metabolites identified by MS/MS and a total of 10 tentatively identified formulae based on exact mass [6]. Dataset 14 contains 10 metabolites identified by MS/MS and 6 tentatively identified formulae based on exact mass [6]. None of these endogenous signals was misclassified as 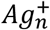-related by our algorithm.

### 4.3. Overlapping peak detection performance

Figure 2 shows a case example where the overlapping peak detection algorithm successfully identified overlapping ions when searching for the *Ag*_6_ cluster in Dataset 1. Figure 2A depicts the experimental mean profile spectrum in the mass range of interest along with the calculated profile of the *Ag*_6_ cluster. While most peaks follow the calculated isotopic distribution, experimental peaks at *m/z* 641.43, *m/z* 643.43 and *m/z* 653.43 are considerably more intense than in the predicted pattern. This generates a mismatch between the experimental and calculated peaks that leads to a low *S*1 score. Figure 2B shows the spatial distributions of each of the ions in the *Ag*_6_ cluster. The correlation map in Figure 2D clearly indicates that peaks at *m/z* 641.43 and *m/z* 643.43 have a spatial distribution that is unlike that of the rest of the ions in the cluster. The peak at *m/z* 653.43 also shows a considerably different spatial correlation to the rest. These low correlations lead to a lower *S*2 score. Figure 2C is a zoom-in of the peaks at *m/z* 641.43 and *m/z* 643.43 showing that the silver ion peaks are clearly overlapped with Ag-unrelated signals.

**Fig. 2.**
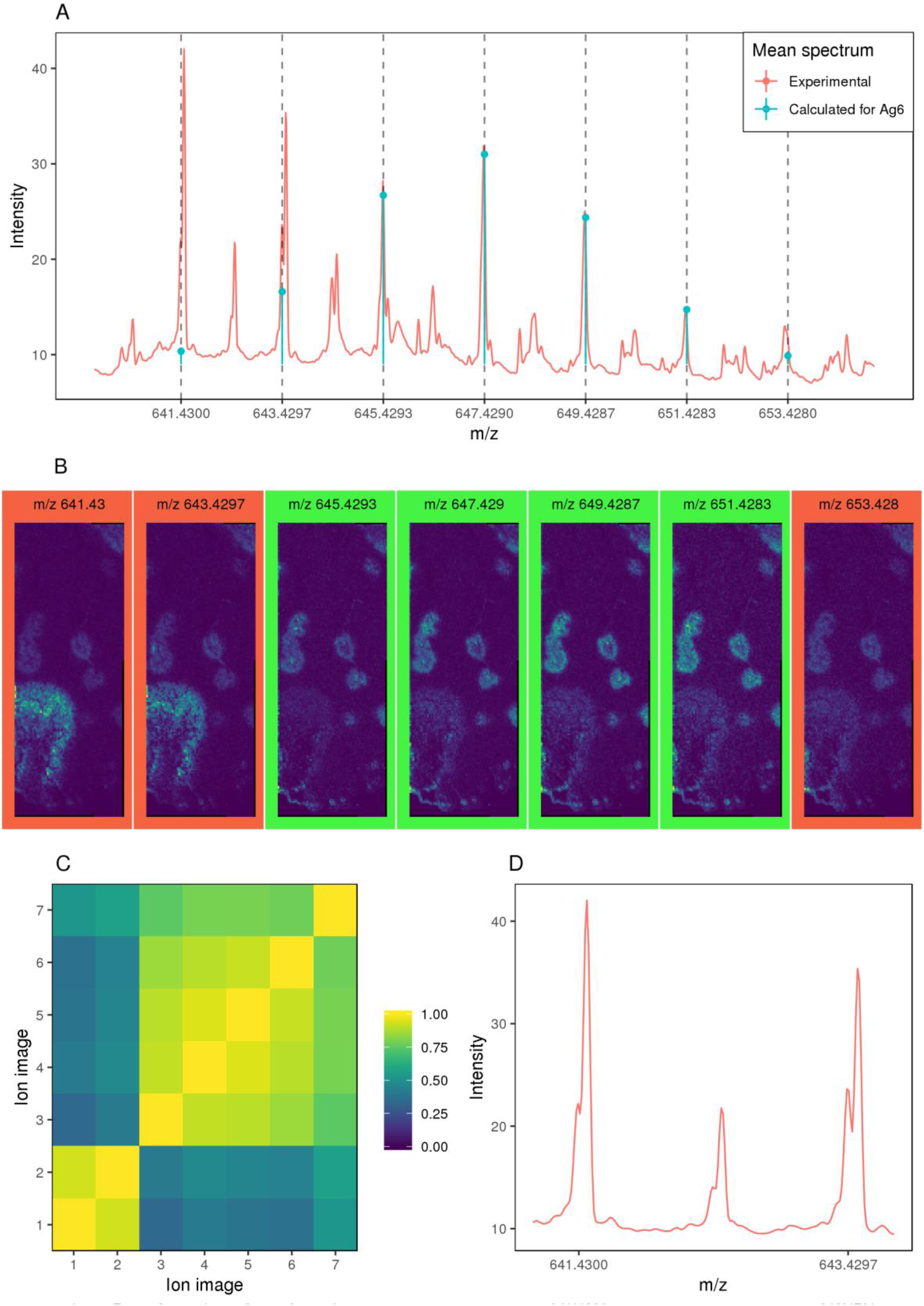
Overlapping detection algorithm performance when searching for the *Ag*_6_ cluster in Dataset 1. (A) Comparison between the mean experimental spectra and the theoretical *Ag*_6_ cluster at the *Ag*_6_ cluster masses within a tolerance of 4 scans. Red and blue represent theoretical and experimental profiles, respectively. As can be seen, while the peaks in the centre of the cluster perfectly match the theoretical ratios, the peaks on the edges differ considerably. (B) Spatial distributions of the experimental cluster peaks. After performing the overlapping detection only the four ion images in the centre in green are classified as Ag-related. The remaining ion images in red are classified as *Ag*^n+^ suffering from overlapping. The morphologies of the *Ag*^n+^ overlapped ions (red) differ from the ones without overlapping (green) due to ion overlapping. (C) Correlation matrix between the experimental ion images of the *Ag*_6_ cluster. The ion image number corresponds to the position of the ion in the isotopic pattern in ascending order of m/z. The first two images are clearly not correlated with the remaining images of the cluster. The last image also shows a considerably lower correlation. (D) Zoom-in of experimental mean spectra. Peaks m/z 641.43 and m/z 643.43 show clear overlapping.

Initially, given the low S1 and S2 scores, all peaks in the *Ag*_6_ cluster were misclassified as not 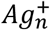-related. Using the overlapping detection algorithm, the peaks at *m/z* 645.43, *m/z* 647.43, *m/z* 649.43 and *m/z* 651.43 were correctly tagged as belonging to *Ag*_6_. Peaks at *m/z* 641.43, *m/z* 643.43 and *m/z* 653.43 were tagged as related to *Ag*_6_ but with overlapping.

Supplementary Figure S9 explores the effects of overlapping peak detection on overall performance. Two main differences can be appreciated. Firstly, there is an overall increase in the *S*_1_ · *S*_2_ score obtained by the “positive class” which leads to a bigger gap between the “positive class” and the “negative class” making the thresholding classification more robust. This is due to the identification of some overlapping peaks in the 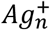 clusters. Additionally, there is a clear improvement in the scores obtained by the *Ag*_6_ cluster. The *Ag*_6_ cluster suffers from overlapping in most of the datasets and is, therefore, the cluster most benefitted from the overlapping detection algorithm. It is also important to note that the overlapping peak detection algorithm does not add any false positives as the *S*_1_ · *S*_2_ remains unchanged for the “negative class”. This proves that overlapping detection leads to less misclassification of 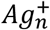-related peaks.

### 4.4. Matrix-related peak annotation improves the post-processing

In order to explore the influence of the annotation and removal of matrix-related peaks in the post-processing workflows, we carried out a multivariate statistical exploratory analysis. The widely used linear algorithm Principal Component Analysis (PCA) [30] was performed on all 14 datasets before and after removal of the 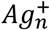 peaks. We then compared the quality of the spatial representation of the first three principal components. Given the lack of a standard quantitative metric to compare the quality of two images in MSI, we followed the trend established by recent work [11,31,32] and performed a qualitative visual comparison.

Figure 3 shows the results of this exploratory analysis on Dataset 2 and Dataset 11. In the pancreatic tissue represented in Figure 3A (Dataset 2), PC1 did not change significantly after matrix removal, while PC2 and PC3 showed a wider variety of morphologies on the tissue after the 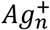 interference was removed. In the brain tissue shown in Figure 3B (Dataset 11) the contrast enhancement is even clearer in the three PCs. Before the 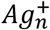 peaks were removed, PC1 and PC3 did not capture any substantial morphology but afterwards, they did and PC2, which already showed morphological information, did so with increased contrast. To convey the three principal components in a single picture we encoded each of them as a colour in the Red Green Blue colour model (RGB). The RGB picture became richer and more informative after the 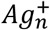 peaks were removed. Similar results were obtained in the remaining 12 datasets and their corresponding images can be accessed in Supplementary Figures S10-S13. The main conclusion that can be drawn from the visual analysis of these results is that the removal of matrix-related peaks leads to a generalized enhancement in the contrast of morphological structures obtained with the first principal components. This is due to the fact that the variance contribution of the matrix-related signals is not fed to the PCA and therefore the resulting principal components are better focused on the morphology of the tissue. In agreement with previous work on the effects of MSI data reduction [33], these results demonstrate that the removal of matrix-related signals improves post-processing, especially when using linear algorithms such as the widely used PCA.

**Fig. 3.**
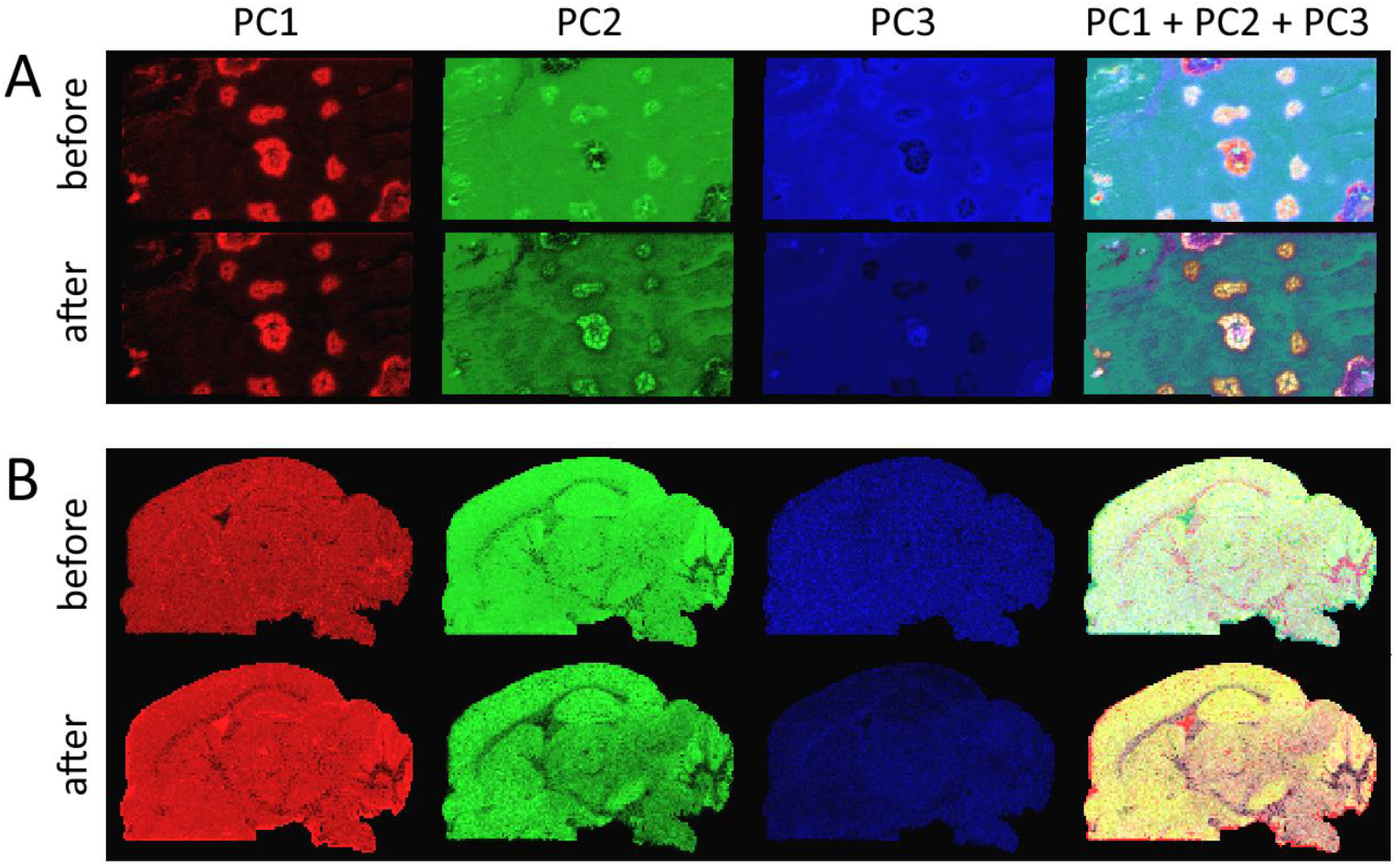
Exploratory analysis with PCA before and after removing matrix-related peaks. Red, green and blue are used to represent the spatial distribution of PC1, PC2 and PC3, respectively. The last column uses the Red Green Blue colour model (RGB) to represent the first three principal components in a single image. The annotation and removal of the matrix-related peaks lead to a generalized improvement in the contrast of morphological structures in all principal components. (A) Pancreas tissue from Dataset 2. (B) Brain tissue from Dataset 11 [18].

## 5. Discussion & Conclusion

The goal of this study was to develop, optimize and validate a new algorithm to annotate signals attributed to the LDI promoting material in MSI. The developed algorithm is packaged and released as rMSIcleanup, an open-source R package freely available for the scientific community and fully integrated with rMSIproc [20], a stand-alone package for the visualization, pre-processing and analysis of MSI datasets.

In comparison to the top-performing alternatives for matrix-related peak annotation which are based on machine and deep learning [11], rMSIcleanup has the main advantage of using two intuitive scores (accounting for the isotopic ratios of clusters and the spatial distribution of their ions) and providing a visual justification of each annotation. This is a key contribution as it helps overcome the black-box problem, increases the user’s confidence in the annotation and can help researchers optimize experimental workflows (for instance, choosing LDI promoters that minimize interferences in the *m/z* range of interest). Another merit of our work is that, to our knowledge, it is the first matrix signal annotation algorithm to explicitly detect and deal with overlapping MS signals, which successfully prevents overlapped peaks from being misclassified. Given that we follow a targeted analytical approach, our classification is focused only on matrix-related signals while the algorithms presented by Ovchinnikova et al. [11] have a broader scope and also classify as off-sample other exogenous compounds. In the era of big data, these two apparently opposite approaches (namely our analytical approach based on chemical similarity scores and their untargeted approach based on machine learning) must not only coexist but also complement each other following the trend already initiated in other fields [34]. This reality urges the MSI community to develop annotation algorithms capable of, not only exploiting the knowledge in the increasingly large amounts of MSI datasets available, but also incorporating metrics that take into account the chemical context of the sample to aid transparent justification.

AgLDI MSI was chosen to validate the algorithm, due to the well-understood ionization of silver. A “validation list” was compiled from the literature, which included silver clusters that should be present in all samples and silver adducts or compounds that should not be present in any of them. Given the heterogeneity of the samples used in this study, the described validation list was adapted to each dataset. For each dataset, those clusters in the validation list for which the experimental data contained none of their theoretical masses were excluded. These adjustments in the validation list prevented an overestimation of the performance of the algorithm attributed to a high number of correctly classified “negative class” clusters (i.e. true negatives) located in mass ranges with no signal. We propose this validation strategy as a novel alternative to more common validation approaches such as chemical standards [6] or expert annotation [11,32]. This study adds to previous work [6,14,17,29,35] and further demonstrates the potentiality of AgLDI MS imaging, a thriving technology known for its reduced background signals in spatial metabolomics that is strongly complemented by our annotation algorithm as it further removes the influence of the matrix.

In agreement with previous work on the effects of MSI data reduction [33], we have demonstrated that the annotation and removal of signals related to the LDI promoting material used can further enhance post-processing, due to the elimination of variables attributed to exogenous compounds that do not reflect the morphology nor chemical composition of the sample. These results highlight the need to include software annotation tools such as rMSIcleanup in MSI workflows before exploring the datasets with classical data analysis techniques used in metabolomics. Here we would like to emphasize the need for a standardized quantitative metric to assess the quality of MSI images and we acknowledge the relevance of standardization initiatives such as the MALDISTAR project (www.maldistar.org).

We envision two main applications for rMSIcleanup. On the one hand, it can be used in a purely exploratory fashion to better understand ionization and adduct cluster formation in new matrices, tissues and applications. In this case, the user is advised to add a long list of potential adducts or neutral losses to assess their formation. The validation approach followed in this paper is a clear example of this exploratory application of rMSIcleanup. A second application is the automated peak annotation of well-known matrices and tissues. In this case, only the clusters that are known to be formed need to be given to the software. This curated selection increases the data-processing speed. The set of matrix-related annotated peaks can then be eliminated from the dataset prior to performing post-processing workflows such as multivariate statistical analysis.

Finally, the promising results obtained in the annotation of 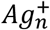-related peaks in AgLDI MSI open the door to the extension of this methodology to more widely used matrices such as 2,5-Dihydroxybenzoic acid (DHB), 1,5-Diaminonaphthalene (DAN), and 9-Aminoacridine (9AA) among others. These organic matrices pose greater challenges. Firstly, they lead to increased matrix background due to their greater fragmentation and adduct formation [36–38] and the higher quantities in which they are added [37]. Moreover, they present the problem of “hot spot” formation given their less homogeneous application process [39]. These issues highlight not only the benefits of AgLDI MSI but also that matrix-related peak annotation can benefit data post-processing even further in applications using organic matrices.

## Author contribution

**Gerard Baquer:** Conceptualization, Methodology, Software, Validation, Visualization, Writing - Original Draft. **Lluc Sementé:** Validation, Writing - Review & Editing. **María Garcia-Altares:** Conceptualization, Methodology, Investigation, Writing - Review & Editing, Supervision. **Young Jin Lee:** Resources, Review & Editing. **Pierre Chaurand:** Resources, Review & Editing. **Xavier Correig:** Conceptualization, Methodology, Writing - Review & Editing, Supervision, Project administration, Funding acquisition. **Pere Ràfols:** Conceptualization, Methodology, Investigation, Writing - Review & Editing, Supervision.

## Acknowledgements

The authors acknowledge the financial support of the Spanish Ministry of Economy and Competitivity through projects TEC2015-69076-P and RTI2018-096061-B-100. GB acknowledges the financial support of the European Union’s Horizon 2020 research and innovation programme under the Marie Skłodowska-Curie grant agreement No. 713679 and the Universitat Rovira i Virgili (URV). LS acknowledges the financial support of Universitat Rovira i Virgili through the pre-doctoral grant 2017PMF-PIPF-60. MGA acknowledges the financial support from the Spanish Ministry of Science, Innovation and Universities (MICIN) through the post-doctoral grant IJCI-2017-33438. PC acknowledges financial support from the Natural Sciences and Engineering Research Council of Canada (NSERC).

