## Supplementary Material for "rMSIcleanup: An open-source tool for matrix-related peak annotation in mass spectrometry imaging and its application to silver-assisted laser desorption/ionization"

### A. Visual Report

```
################################################################################
– Package Version: 0.1.1
– Time: 2019–07–23 13:39:26
################################################################################
IMAGE INFORMATION
– Peak matrix: 2016_06_01_Brain_Control–Ag.tar
– Full spectrum: NULL
– Number of peaks: 135
– Number of pixels: 10010
– Mass Range: [ 81.0431355625551 , 1002.15406104405 ]
################################################################################
MATRIX INFORMATION
– Matrix formula: Ag1; Ag1F1; Ag1Cl1; Ag1Br1; Ag1I1; Ag1H1; Ag1H2; Ag1He1; Ag1N1O3; Ag1Th2; Ag1F2; Ag1B1F4; Ag1C27H
Ag1C26H54O1; Ag1C28H58O1; Ag1C30H62O1; Ag1C26H52O2; Ag1C30H60O2
– Add list: F1; Cl1; Br1; I1; H1; H2; He1; N1O3; Th2; F2; B1F4; C27H56; C29H60; C31H64; C26H54O1; C28H58O1; C30H62O1;
– Substract list: NULL
– Maximum cluster multiplication: 10
– Base forms: Ag1; Ag1H1; Ag1H2; Ag1He1; Ag1F1; Ag1Cl1; Ag1F2; Ag1N1O3; Ag1Br1; Ag1B1F4; Ag2; Ag2H2; Ag2H4; Ag2He2
Ag2F4; Ag3; Ag3H3; Ag3H6; Ag3He3; Ag2N2O6; Ag2Br2; Ag3F3; Ag2B2F8; Ag3Cl3; Ag4; Ag4H4; Ag3F6; Ag4H8; Ag4He4; Ag2I2
Ag1C26H52O2; Ag4F4; Ag3N3O9; Ag1C29H60; Ag1C28H58O1; Ag5; Ag5H5; Ag1C31H64; Ag5H10; Ag1C30H62O1; Ag5He5; Ag
Ag4F8; Ag3B3F12; Ag5F5; Ag6; Ag6H6; Ag6H12; Ag6He6; Ag4N4O12; Ag3I3; Ag5Cl5; Ag5F10; Ag4Br4; Ag7; Ag7H7; Ag6F6; Ag
Ag5N5O15; Ag6Cl6; Ag8; Ag8H8; Ag6F12; Ag8H16; Ag7F7; Ag8He8; Ag5Br5; Ag4I4; Ag9; Ag5B5F20; Ag9H9; Ag2C54H112; Ag2
Ag9He9; Ag2C52H104O4; Ag8F8; Ag6N6O18; Ag7F14; Ag2C58H120; Ag2C56H116O2; Ag10; Ag10H10; Ag2C62H128; Ag10H20
Ag2C60H120O4; Ag9F9; Ag8Cl8; Ag2Th4; Ag8F16; Ag6B6F24; Ag5I5; Ag7N7O21; Ag10F10; Ag9Cl9; Ag7Br7; Ag9F18; Ag8N8O2
Ag10F20; Ag3C81H168; Ag3C78H162O3; Ag8Br8; Ag3C78H156O6; Ag9N9O27; Ag3C87H180; Ag8B8F32; Ag3C84H174O3; Ag3
Ag3C90H180O6; Ag10N10O30; Ag3Th6; Ag9B9F36; Ag10Br10; Ag8I8; Ag10B10F40; Ag4C108H224; Ag4C104H216O4; Ag4C104
Ag9I9; Ag4C124H256; Ag4C120H248O4; Ag4C120H240O8; Ag4Th8; Ag10I10; Ag5C135H280; Ag5C130H270O5; Ag5C130H260
Ag5C155H320; Ag5C150H310O5; Ag5C150H300O10; Ag5Th10; Ag6C162H336; Ag6C156H324O6; Ag6C156H312O12; Ag6C174
Ag6C180H372O6; Ag6C180H360O12; Ag7C189H392; Ag7C182H378O7; Ag6Th12; Ag7C182H364O14; Ag7C203H420; Ag7C196
Ag8C216H448; Ag8C208H432O8; Ag7C210H420O14; Ag7Th14; Ag8C208H416O16; Ag8C232H480; Ag8C224H464O8; Ag8C248
Ag9C234H486O9; Ag8C240H480O16; Ag9C234H468O18; Ag8Th16; Ag9C261H540; Ag9C252H522O9; Ag10C270H560; Ag9C27
Ag10C260H520O20; Ag9C270H540O18; Ag9Th18; Ag10C290H600; Ag10C280H580O10; Ag10C310H640; Ag10C300H620O10;
################################################################################
PROCESSING INFORMATION
– S1 threshold: 0.8
– S2 threshold: 0.8
– Similarity method: euclidean
– Magnitude of interest: intensity
– Tolerance mode: scans
– Tolerance scans: 4
################################################################################
```

*Fig. S1 Initial summary of the results including main metrics of the images, the chemical formulae, the potential cluster adduct and neutral losses.*

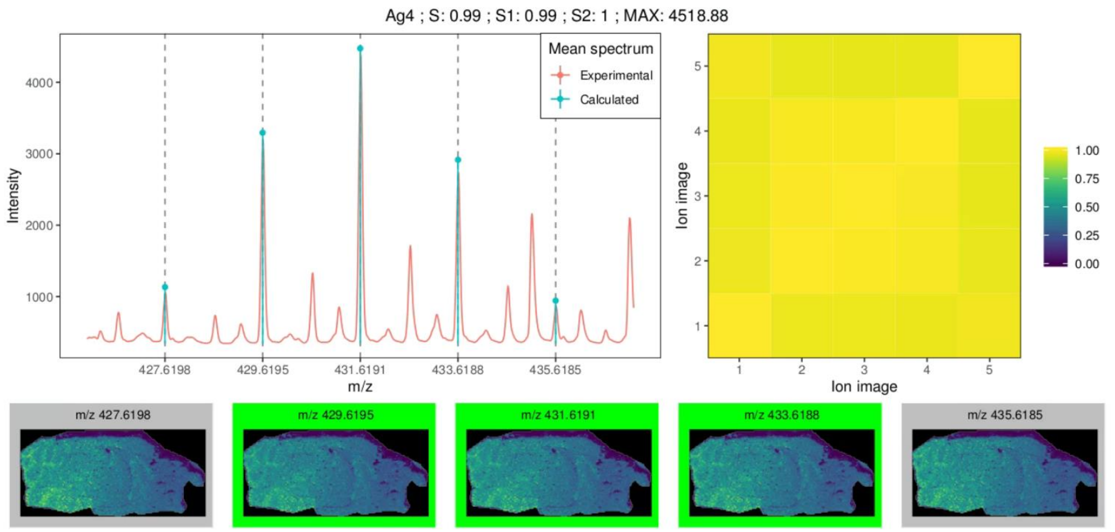


*Fig. S2 Visual report for cluster Ag4 in Dataset 7. The report includes the comparison between experimental and calculated peaks, the correlation map and all ionic images. The ionic images with a green border are tagged as silver-related. The ionic images with a grey border are not found in the peak list provided and are thus not classified.*

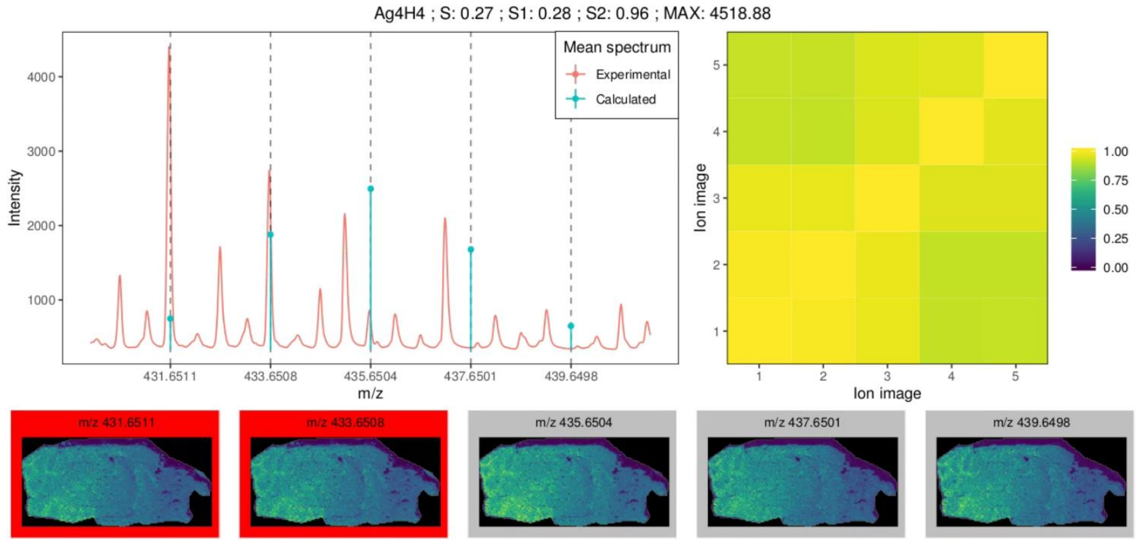


*Fig. S3 Visual report for cluster Ag4H4 in Dataset 7. The report includes the comparison between experimental and calculated peaks, the correlation map and all ionic images. The ionic images with a red border are tagged as not silver-related. The ionic images with a grey border are not found in the peak list provided and are thus not classified.*

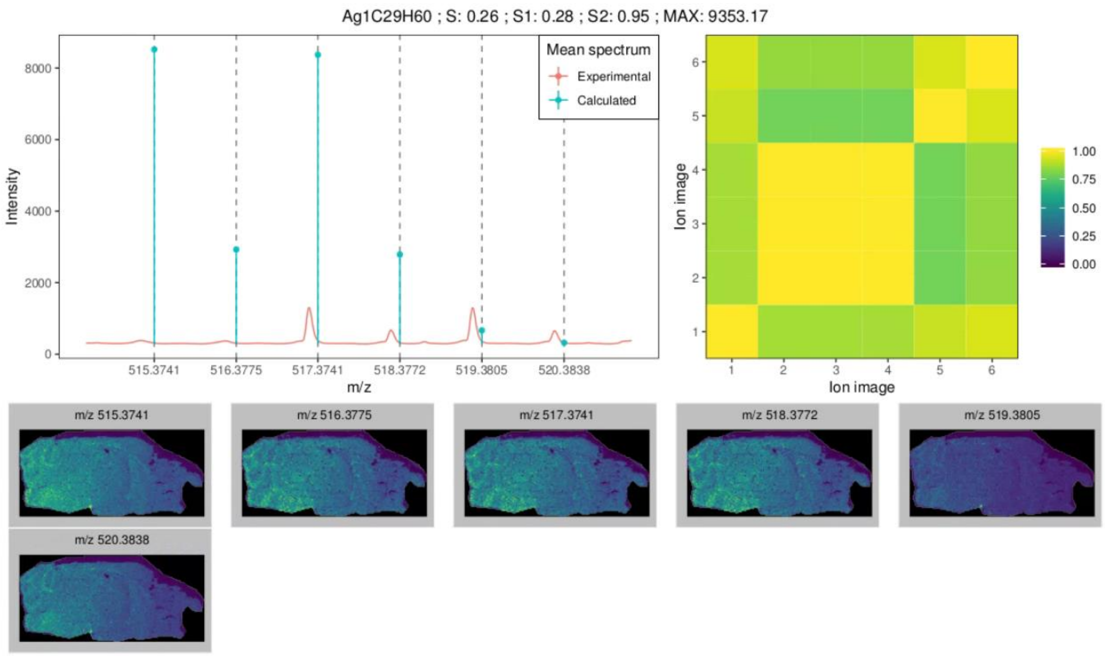


*Fig. S4 Visual report for cluster AgC29H60 in Dataset 7. The report includes the comparison between experimental and calculated peaks, the correlation map and all ionic images. The ionic images with a grey border are not found in the peak list provided and are thus not classified.*

### B. Table of cluster numbers

| | | | | | |
|---|---|---|---|---|---|
| 1 | $Ag_1$ | 28 | $Ag_4Cl_4$ | 55 | $Ag_7F_7$ |
| 2 | $Ag_2$ | 29 | $Ag_6He_6$ | 56 | $C_{60}H_{124}O_2 + Ag_2$ |
| 3 | $Ag_5$ | 30 | $Ag_3Br_3$ | 57 | $Ag_5B_5F_{20}$ |
| 4 | $Ag_7$ | 31 | $Ag_4H_4$ | 58 | $Ag_8Cl_8$ |
| 5 | $Ag_4$ | 32 | $Ag_1N_1O_3$ | 59 | $Ag_8H_8$ |
| 6 | $Ag_9$ | 33 | $C_{26}H_{52}O_2 + Ag$ | 60 | $C_{26}H_{54}O_1 + Ag_1$ |
| 7 | $Ag_3$ | 34 | $C_{29}H_{60} + Ag$ | 61 | $C_{60}H_{120}O_4 + Ag_2$ |
| 8 | $C_{28}H_{58}O_1 + Ag$ | 35 | $Ag_5F_{10}$ | 62 | $Ag_9H_9$ |
| 9 | $Ag_8$ | 36 | $Ag_2H_4$ | 63 | $Ag_4B_4F_{16}$ |
| 10 | $Ag_6$ | 37 | $Ag_5Cl_5$ | 64 | $Ag_2F_4$ |
| 11 | $Ag_3Cl_3$ | 38 | $Ag_1I_1$ | 65 | $Ag_5He_5$ |
| 12 | $Ag_2TH_4$ | 39 | $Ag_9He_9$ | 66 | $Ag_7N_7O_{21}$ |
| 13 | $Ag_2H_2$ | 40 | $Ag_6F_{12}$ | 67 | $Ag_9F_9$ |
| 14 | $C_{30}H_{60}O_2 + Ag$ | 41 | $Ag_1Cl_1$ | 68 | $Ag_8F_{16}$ |
| 15 | $Ag_6F_6$ | 42 | $C_{54}H_{112} + Ag_2$ | 69 | $Ag_{10}He_{10}$ |
| 16 | $Ag_7H_7$ | 43 | $Ag_3H_6$ | 70 | $Ag_4N_4O_{12}$ |
| 17 | $Ag_1F_2$ | 44 | $Ag_7He_7$ | 71 | $Ag_6B_6F_{24}$ |
| 18 | $Ag_3I_3$ | 45 | $C_{52}H_{108}O_2 + Ag_2$ | 72 | $Ag_7F_{14}$ |
| 19 | $Ag_1H_2$ | 46 | $Ag_6H_{12}$ | 73 | $C_{52}H_{104}O_4 + Ag_2$ |
| 20 | $Ag_5F_5$ | 47 | $C_{62}H_{128} + Ag_2$ | 74 | $C_{56}H_{116}O_2 + Ag_2$ |
| 21 | $Ag_1He_1$ | 48 | $Ag_1B_1F_4$ | 75 | $Ag_9H_{18}$ |
| 22 | $Ag_6Cl_6$ | 49 | $Ag_6H_6$ | 76 | $Ag_8F_8$ |
| 23 | $Ag_{10}$ | 50 | $Ag_7H_{14}$ | 77 | $C_{58}H_{120} + Ag_2$ |
| 24 | $Ag_4Br_4$ | 51 | $Ag_5N_5O_{15}$ | 78 | $Ag_6Br_6$ |
| 25 | $Ag_3F_6$ | 52 | $Ag_5I_5$ | 79 | $Ag_{10}H_{10}$ |
| 26 | $Ag_4I_4$ | 53 | $Ag_5H_{10}$ | 80 | $Ag_{10}H_{20}$ |
| 27 | $Ag_8He_8$ | 54 | $Ag_8H_{16}$ | 81 | $Ag_7Cl_7$ |

*Table S1. Cluster numbers used in Figure 1C in decreasing order of mean S1·S2 performance*

### C. Example clusters

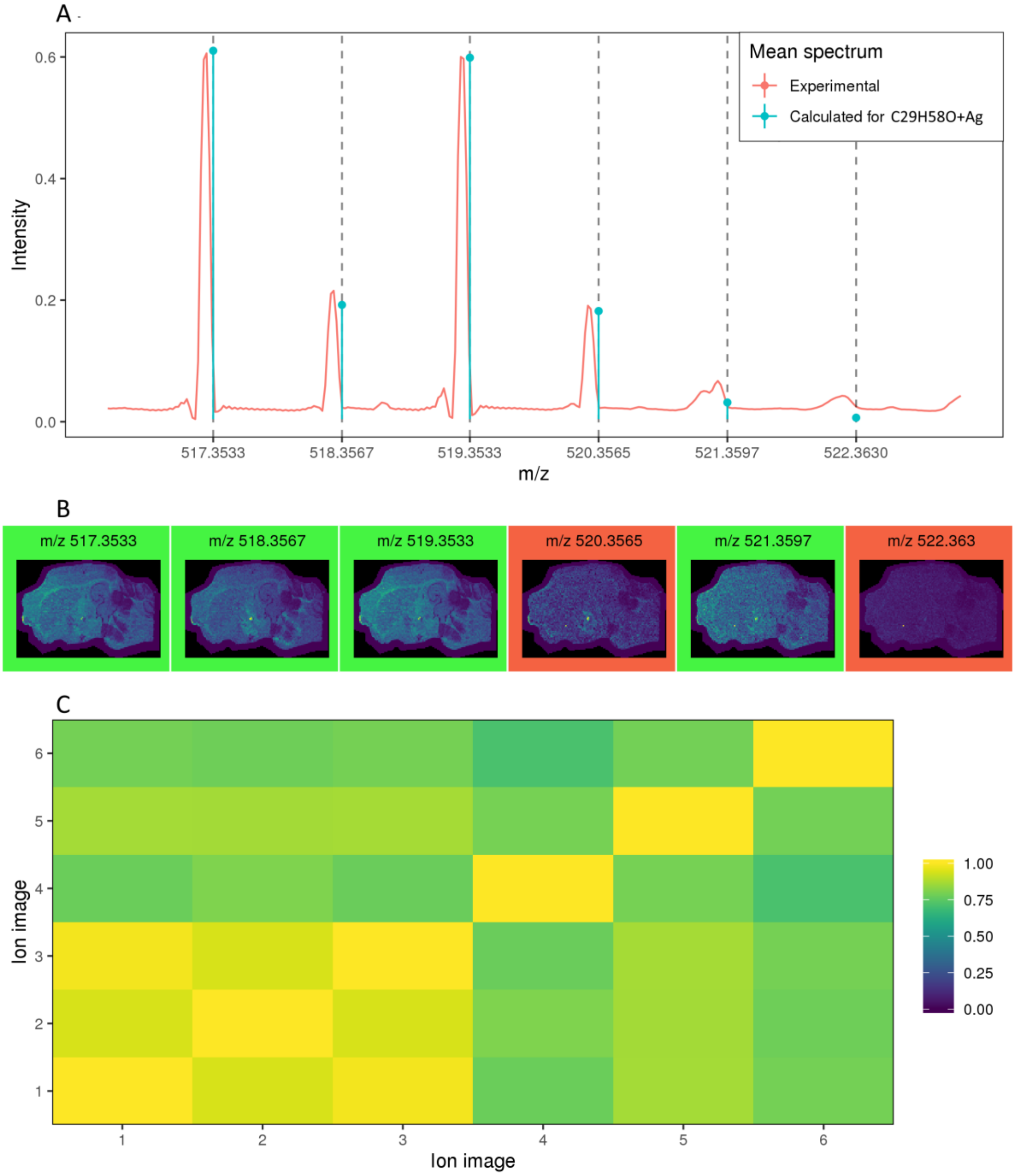


*Fig. S5 Classification results of cluster $C_{28}H_{58}O + Ag$ in Dataset 4. (A) Comparison between the mean experimental spectra and the theoretical pattern. (B) Spatial distributions of the experimental cluster peaks. (C) Correlation matrix between the experimental ionic images of the cluster. The cluster is misclassified as silver-related (false positive). Further study and annotation of these peaks would be needed to assess if the compound is indeed present in the sample implying that this specific compound should not be included in the "ground truth" as a negative class. Nevertheless, the constant and notable mass error between experimental and theoretical peaks allows us to infer that the experimental pattern might be due to a different compound. Adjusting the mass tolerance of the algorithm would get rid of these false positives.*

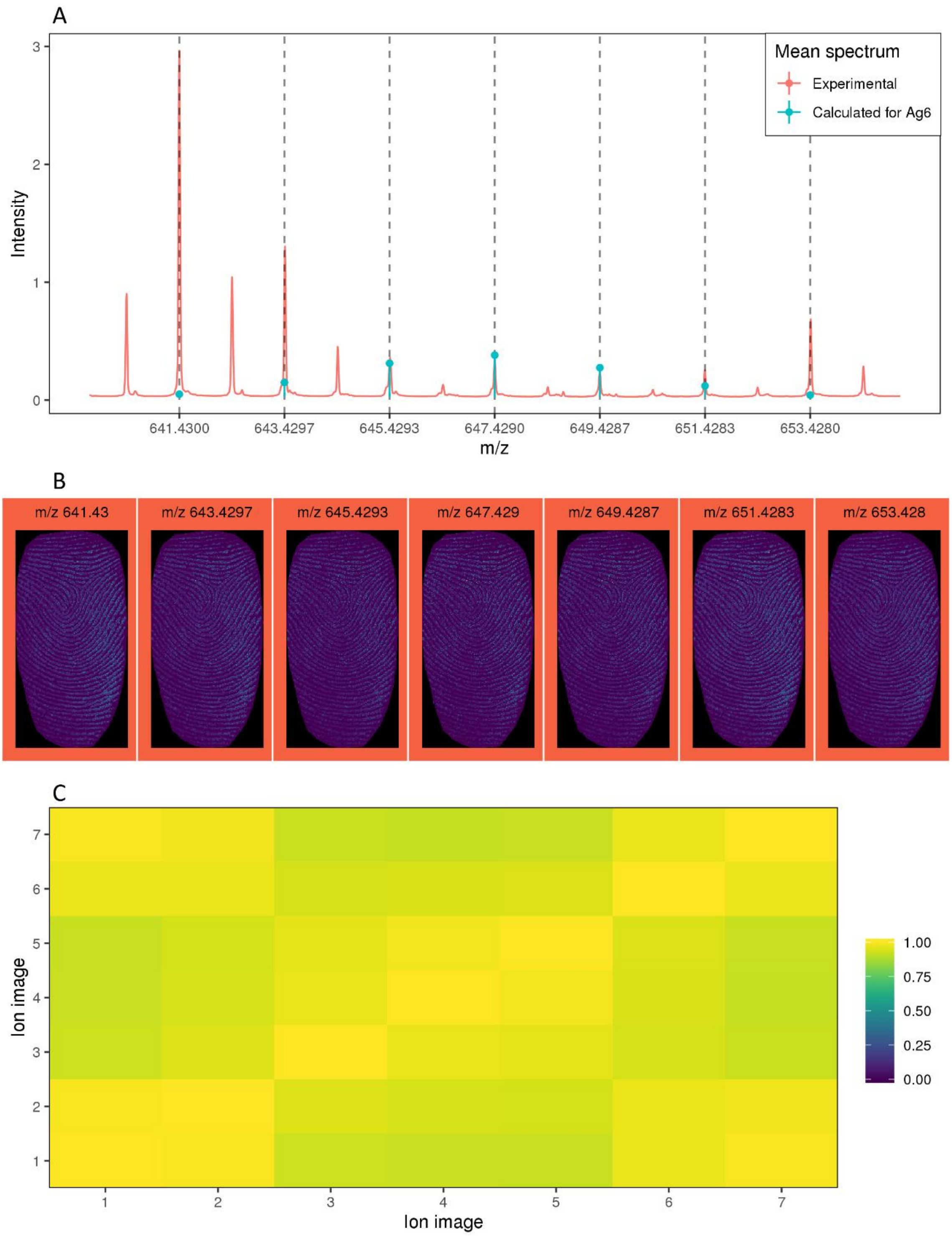


*Fig. S6 Classification results of cluster $Ag_6$ in Dataset 12. (A) Comparison between the mean experimental spectra and the theoretical pattern. (B) Spatial distributions of the experimental cluster peaks. (C) Correlation matrix between the experimental ionic images of the cluster. The cluster is misclassified as not silver-related (false negative). Like the example in Figure 2, peaks m/z 641.43, m/z 643.43 and m/z 653.43 clearly suffer from overlapping. Nevertheless, due to the high homogeneity of the fingerprint sample, the morphological correlation between the overlapped and the non-overlapped ions is relatively high. The overlapping detection algorithm fails to detect the overlapped peaks.*

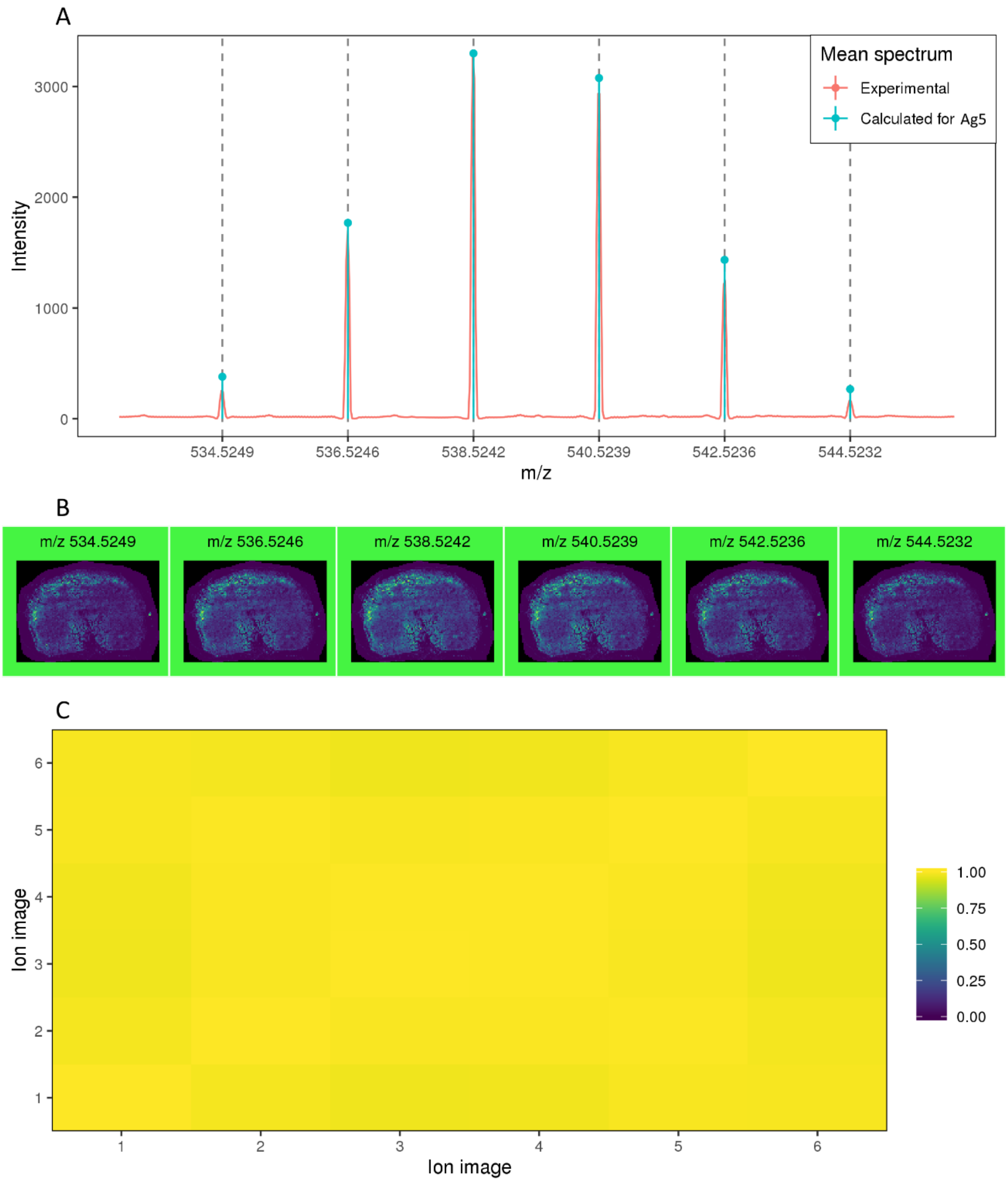


*Fig. S7 Classification results of cluster $Ag_5$ in Dataset 3. (A) Comparison between the mean experimental spectra and the theoretical pattern. (B) Spatial distributions of the experimental cluster peaks. (C) Correlation matrix between the experimental ionic images of the cluster. The cluster is correctly classified as silver-related (true positive).*

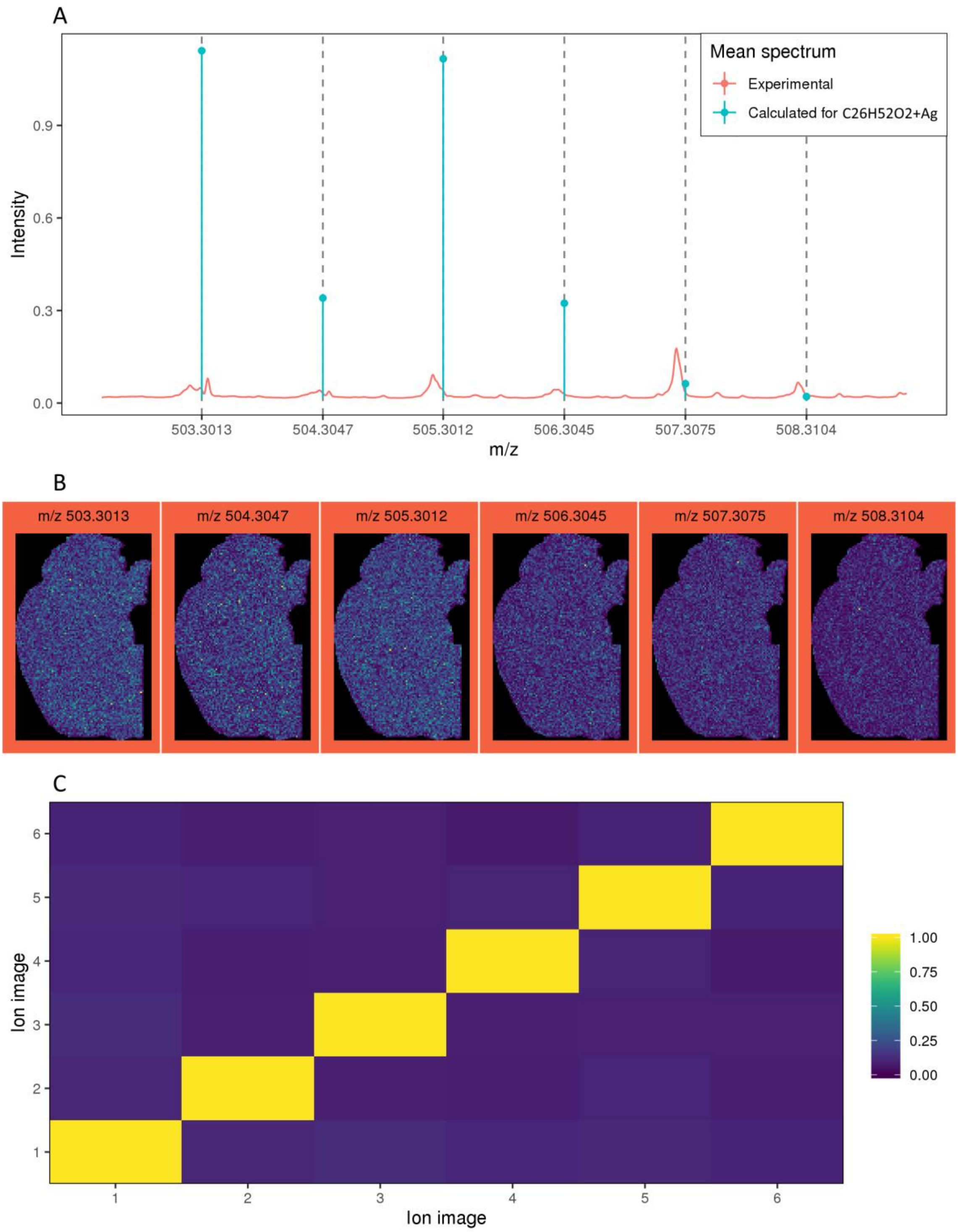


*Fig. S8 Classification results of cluster $C_{26}H_{52}O_2 + Ag$ in Dataset 11. (A) Comparison between the mean experimental spectra and the theoretical pattern. (B) Spatial distributions of the experimental cluster peaks. (C) Correlation matrix between the experimental ionic images of the cluster. The cluster is correctly classified as not silver-related (false positive).*

### D. Effects of overlapping peak detection

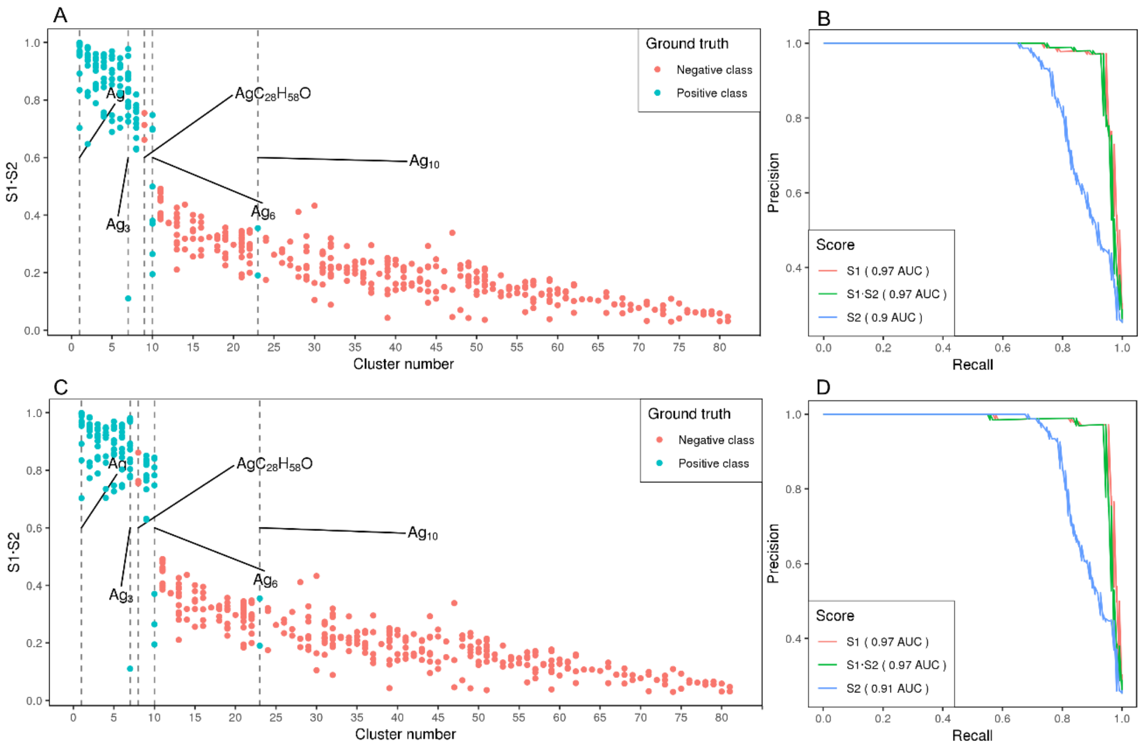


*Fig. S9 Similarity score S1·S2 vs. Cluster number and Precision vs. Recall curves with overlapping peak detection disabled or enabled. (A) & (B) Overlapping peak detection disabled. Multiple Ag6 clusters receive a low score and are thus misclassified as not Ag-related. (C) & (D) Overlapping peak detection enabled. The number of misclassified Ag6 clusters is considerably reduced. Additionally, the gap between the positive and negative class is now bigger leading to a more robust thresholding.*

### E. Complete exploratory analysis using PCA

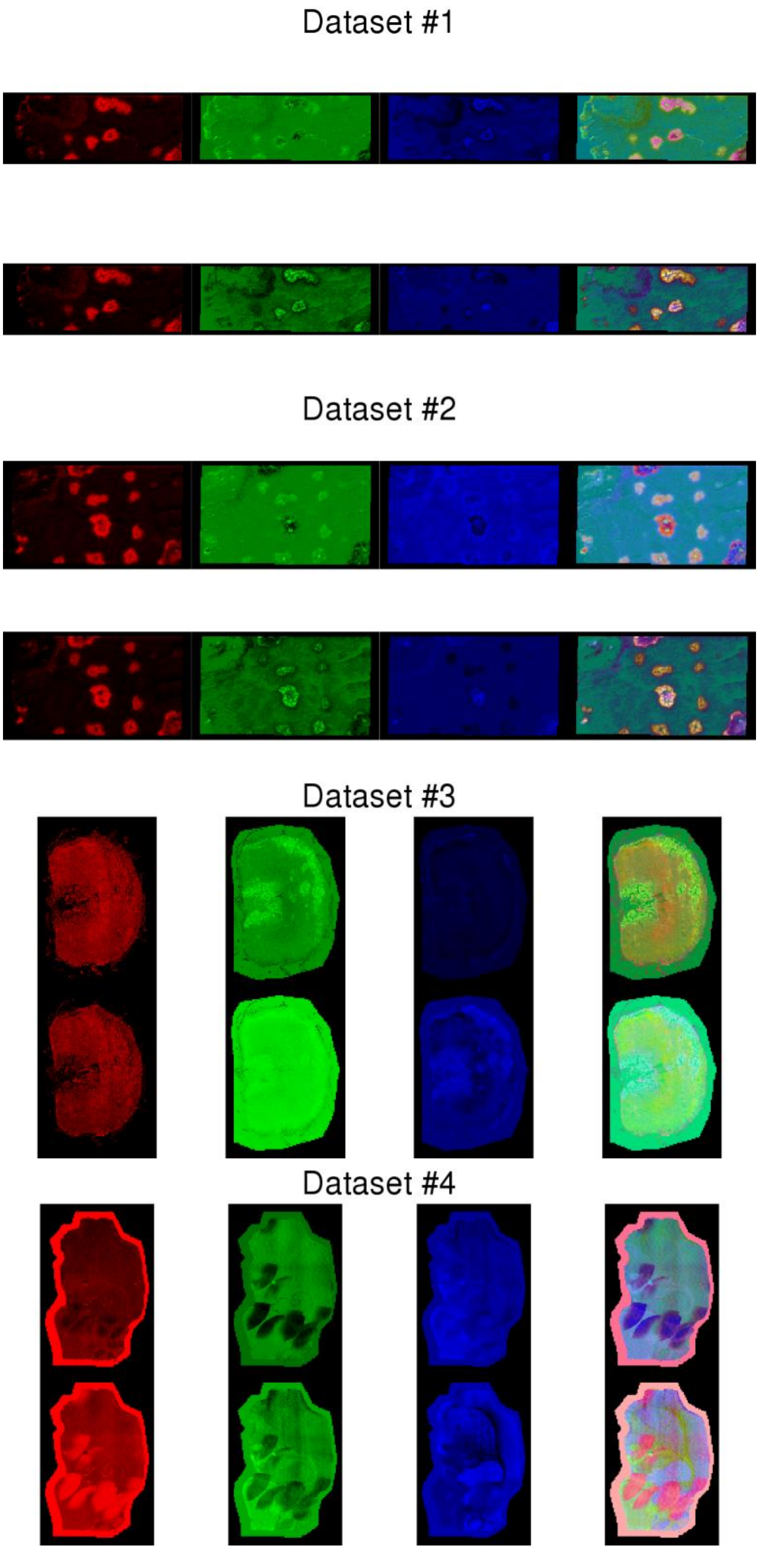


*Fig. S10 Exploratory analysis with PCA before (top row) and after (bottom row) removing matrix-related peaks for Datasets 1-4. Red, green and blue are used to represent the spatial distribution of PC1, PC2 and PC3, respectively. The last column uses the Red Green Blue colour model (RGB) to represent the first three principal components in a single image.*

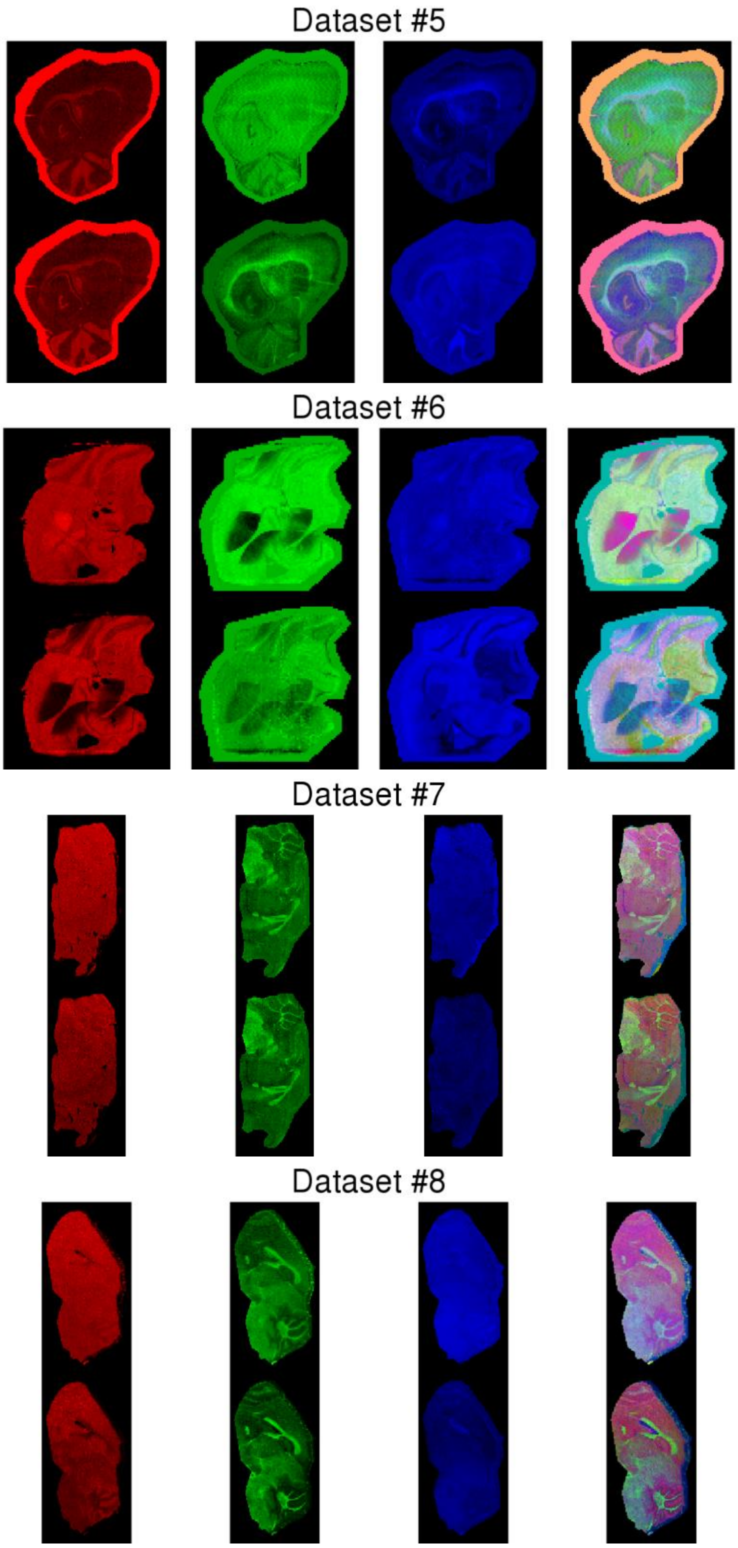


*Fig. S11 Exploratory analysis with PCA before (top row) and after (bottom row) removing matrix-related peaks for Datasets 5-8. Red, green and blue are used to represent the spatial distribution of PC1, PC2 and PC3, respectively. The last column uses the Red Green Blue colour model (RGB) to represent the first three principal components in a single image.*

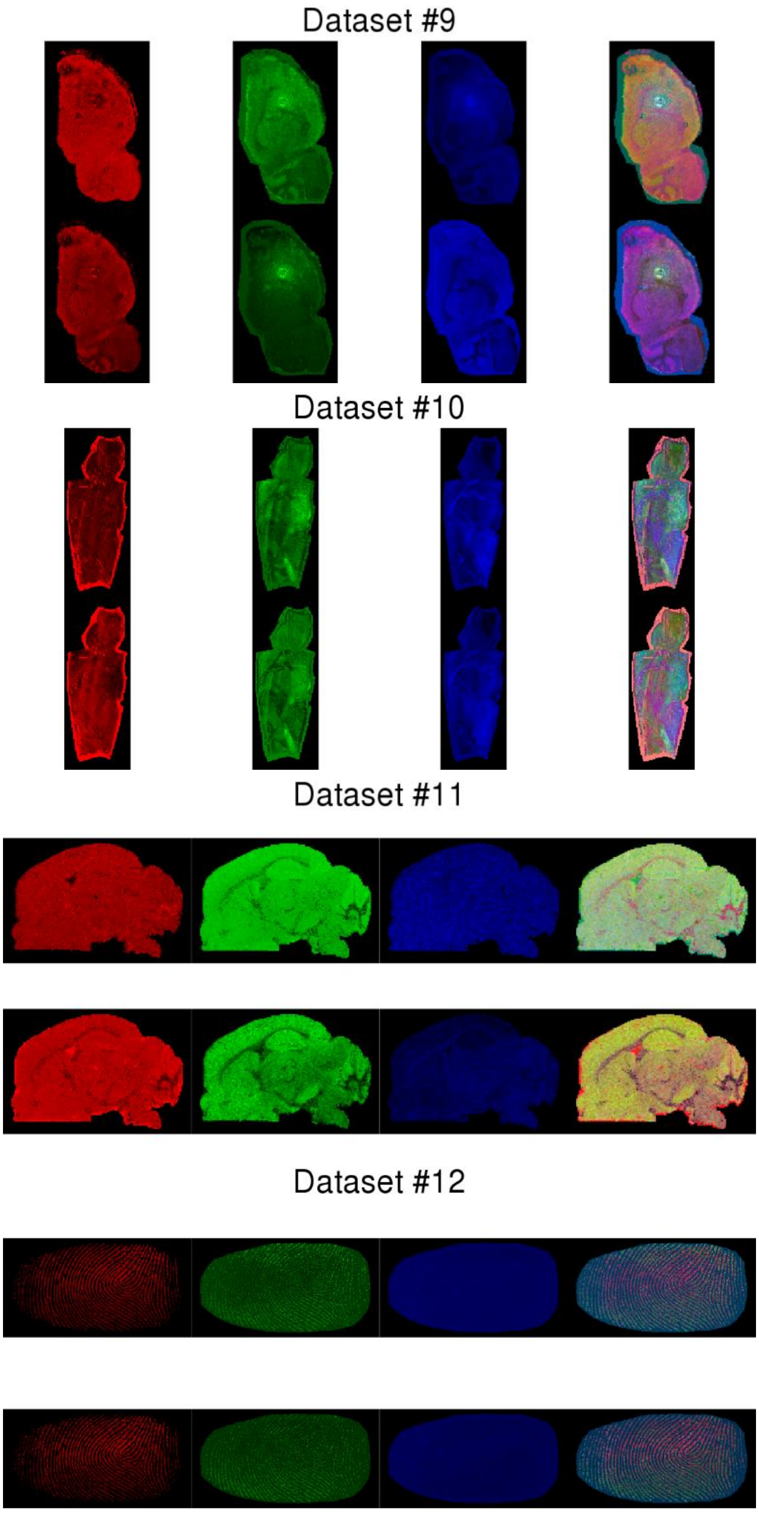


*Fig. S12 Exploratory analysis with PCA before (top row) and after (bottom row) removing matrix-related peaks for Datasets 9-12. Red, green and blue are used to represent the spatial distribution of PC1, PC2 and PC3, respectively. The last column uses the Red Green Blue colour model (RGB) to represent the first three principal components in a single image.*

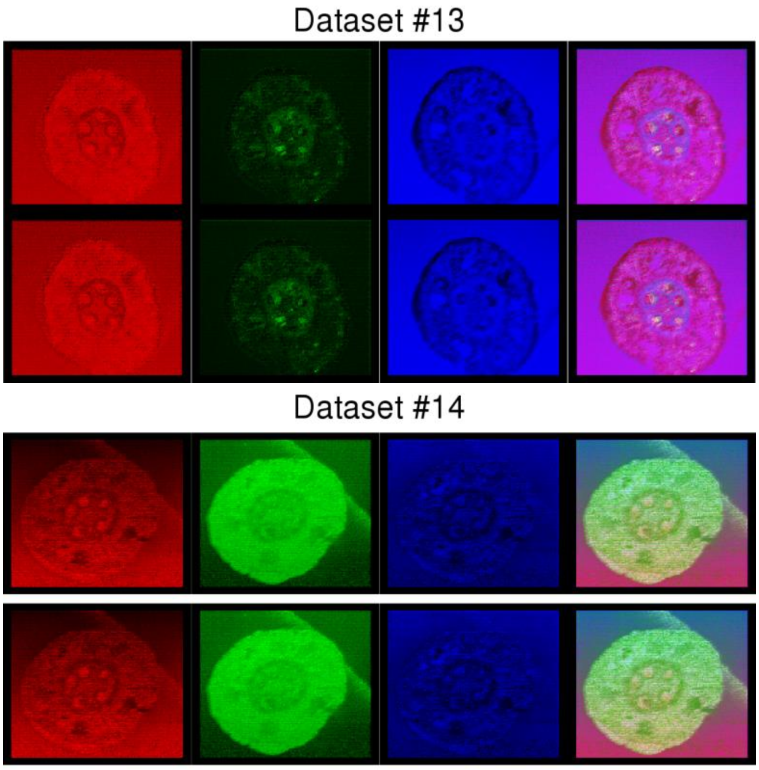


*Fig. S13 Exploratory analysis with PCA before (top row) and after (bottom row) removing matrix-related peaks for Datasets 13-14. Red, green and blue are used to represent the spatial distribution of PC1, PC2 and PC3, respectively. The last column uses the Red Green Blue colour model (RGB) to represent the first three principal components in a single image.*
